# Obesity-driven changes in ECM composition promote local invasion and metastasis of breast tumors

**DOI:** 10.1101/2020.01.29.924431

**Authors:** Andrew Wishart, Yifan Peng, Sydney Conner, Justinne Guarin, Rebecca Crews, Jackson Fatherree, Andrew S. Greenberg, Madeleine J. Oudin

## Abstract

The extracellular matrix (ECM) is a major component of the tumor microenvironment that supports cellular growth, promotes local invasion from the primary tumor, and contributes to metastatic outgrowth in sites of colonization. Obesity is a systemic disease that causes chronic inflammation which can lead to ECM deposition and ultimately fibrosis in adipose tissues such as the mammary gland. Overweight breast cancer patients have increased metastasis to the lung and liver, exhibit resistance to chemotherapy and have worse outcomes. We found that ECM isolated from the mammary gland of both tumor-bearing and obese mice increased invasion of breast cancer cells and set out to investigate whether obesity-driven changes in ECM could identify novel drivers of invasion and metastasis in breast cancer. We performed proteomics of the mammary fat pads of both lean and obese mice and identified the entire landscape of obesity-driven ECM changes. In particular, we focused on Collagen VI, an ECM protein secreted by adipocytes in mammary tissues. Collagen VI is upregulated in the ECM of obese and tumor-bearing mice and is associated with poor outcome in human breast cancer. We found that Collagen VI drives adhesion, migration and invasion of several human breast cancer cell lines via crosstalk between the adhesion receptor NG2 and the receptor tyrosine kinase EGFR, and activation of MAPK signaling. Overall, these studies demonstrate that obesity can have profound effects on the ECM composition of tissues, which in turn can promote local invasion and metastasis.

## Introduction

Obesity is a systemic disease that affects the composition and physiological properties of the brain and peripheral tissues such as the gut, lung, liver, pancreas and adipose tissue. Obesity is a global pandemic: according to the CDC, 70% of US adults are overweight or obese, with the highest prevalence amongst Blacks and Hispanics^1^. Importantly, obesity contributes to 20% of cancer-related deaths, with a higher incidence of breast cancer in obese individuals. Epidemiological studies reveal that the risk of triple-negative breast cancer (TNBC) is associated with an increase in Body Mass Index (BMI) and that a higher proportion of obese patients suffered from TNBC^2–8^. Furthermore, individuals with a high BMI have increased rates of metastasis and decreased response to chemotherapy^6^. TNBC, which represents approximately 25% of breast cancer cases, remains extremely hard to treat, due to its heterogeneous nature and the lack of driver mutations which can be directly targeted^9^. TNBC also has a recurrence rate of over 30%, which is highest in the first three years after diagnosis^10^. Given the increasing rates of obesity and the poor outcomes associated with TNBC, it is important to gain a detailed understanding of the mechanisms of obesity-driven breast cancer progression.

Obesity is thought to drive cancer cell proliferation due to increased hormone, growth factor and cytokine secretion by adipocytes and/or macrophages^11^. The majority of existing literature investigates mechanisms of obesity-induced adipocyte secretion of cytokines which promote inflammation and metabolic substrates such as fatty acids, glucose, and glycerol, which promote tumor cell proliferation proliferation^12^. However, obesity is also associated with fibrosis, the modification of both the amount and composition of extracellular matrix (ECM) proteins. Obesity-associated macrophages promote myofibroblast activation, leading to ECM deposition and ultimately fibrosis in adipose tissue^13^. As a consequence of these complex physiological changes, the local microenvironment of tumors in obese patients is drastically different than in lean patients, suggesting that predicting and targeting cancer metastasis in obese patients requires a more in depth understanding of the obese tumor microenvironment and how it contributes to local invasion. A recent study explored the role of the ECM in obesity-induced breast cancer invasion^14^. Adipose stromal cells isolated from mammary fat pads of obese and lean mice were plated *in vitro* for 2 weeks, where they secreted their own ECM. ECM generated from cells from obese mice showed increased abundance of fibronectin and collagen crosslinking, leading to increased breast cancer cell invasion via mechano-signaling. These studies demonstrated for the first time that obesity impacts the ability of stromal cells isolated from the mammary gland to secrete ECM *in vitro*. However, these experiments focused on ECM generated *in vitro* and did not characterize the landscape of ECM changes associated with obesity.

We and others have shown that the extracellular matrix (ECM) is a major driver of local invasion and metastasis in breast cancer^15,16^. Recent advances in the field of ECM proteomics have increased our understanding of the complexity and heterogeneity of the breast tumor matrisome^16,17^. Mayorca-Giuliani *et al.* describe the matrisome of breast tumor tissue generated via syngeneic 4T1 xenografts relative to normal mammary gland^18^, while Naba *et al* characterize the ECM of highly vs. poorly metastatic breast tumor ECM using LM2 and MDA-MB-231 cells in immunocompromised mice^19^. While these studies revealed the presence of well-studied ECM proteins such as Fibronectin (FN) and Collagen I and identified novel drivers of metastasis, they also highlighted the diversity of individual cues in ECM proteins in tumors, many of which whose function has been poorly studied until now. Here, we characterized the matrisome of the obese mouse mammary gland and show that obese ECM promotes breast cancer cell invasion. We identify Collagen VI, an ECM protein secreted by breast adipocytes^20^ upregulated in tumor and obese mammary ECM, as a novel driver of invasion and metastasis in breast cancer. Overall, these studies provide a novel mechanism by which obesity contributes to tumor progression and metastasis.

## Methods

### Reagents, antibodies, ECM substrates, growth factors, peptides, inhibitors

Reagents were purchased from Fisher Scientific (Hampton, NH) or SIGMA (St. Louis, MO) unless otherwise specified. Antibodies used were: anti-IgG2a isotype control (ab18414; Abcam, Cambridge, MA), anti-Collagen VI (ab199720; Abcam, Cambridge, MA), anti-Fibronectin (ab2413; Abcam, Cambridge, MA), anti-Integrin β1 (12594-1-AP; Proteintech, Rosemont, IL), anti-α-Tubulin (T9026; SIGMA, St. Louis, MO), anti-Histone H3 (ab1791; Abcam, Cambridge, MA), anti-GAPDH (14C10; Cell Signaling Technology, Danvers, MA), anti-phospho-p44/42 ERK (Thr202/Tyr204) (4370; Cell Signaling Technology, Danvers, MA), anti-pFAK397 (3283; Cell Signaling Technology, Danvers, MA). ECM substrates used were: Collagen VI (ab7538; Abcam, Cambridge, MA), Collagen I (CB-40236; Fisher Scientific, Hampton, NH), Fibronectin p (F1141; SIGMA, St. Louis, MO), Elastin (E7277; SIGMA, St. Louis, MO), Laminin (L6274; SIGMA, St. Louis, MO). Endotrophin (ETP) peptide, a 77 amino acid length peptide with >98% purity as determined by analytical HPLC was synthesized by Thermo Fisher Scientific (Waltham, MA). Inhibitors used were: Anti-NG2 (NG2ab) (LS-C22113; Lifespan Biosciences, Seattle, WA), anti-Integrin β1 (β1ab) (P4C10; MilliporeSIGMA, Burlington, MA), Lapatinib (S2111; Selleckchem, Houston, TX), Alpelisib (HY-15244; MedChemExpress, Monmouth Junction, NJ), Defactinib (VS-6063; Selleckchem, Houston, TX), Trametinib (S2673; Selleckchem, Houston, TX).

### Cell culture

MDA-MB-231, MDA-MB-468 and MCF-10A cells were obtained from ATCC (Manassas, VA). MDA-MB-231 and MDA-MB-468 cells were cultured in DMEM with 10% serum and Pen-Strep Glutamine, while MCF-10A cells were cultured as per ATCC recommendations in MEGM bullet kit growth media (CC-3150; Lonza, Walkersville, MD) without gentamycin-amphotericin B mix but with additional 100ng/ml cholera toxin (C8052; SIGMA, St. Louis, MO). Cells were routinely checked for the presence of mycoplasma by a PCR based method using a Universal Mycoplasma Detection Kit (30-1012K; ATCC, Manassas, VA). Only mycoplasma negative cells were used in this study.

### Animal experiments

We used two mouse models of metastatic breast cancer. For xenograft tumors, MDA-MB-231 cells (2 million per mouse in PBS and 20% collagen I) were injected into the fourth right mammary fat pad of three 6-wk-old female NOD-SCID mice (Taconic, Hudson, NY). Once the tumors had all reached 1cm diameter mice were euthanized by CO_2_ and tumors excised. PyMT-MMTV mice (JAX) were used for a genetic mouse model of breast cancer. Female mice were left to grow for 12 weeks until tumor burden reached 2cm^3^. Obese mice were generated with a diet. Female C57BL/6 mice were fed on chow diet Teklad 2016S or high fat diet (Research diets D12492, 60% kcal% fat) for 18 weeks where after mice were euthanized by CO_2_ and mammary fat pads excised.

### Decellularization and protein isolation

Tumors dissected from NOD-SCID mice and PyMT-MMTV mice and mammary fat pads from C57BL/6 mice and were submerged in 0.1% SDS solution and stirred for several days, replacing the solution twice daily. Once the tissue turned completely white (visual sign of successful decellularization) the solution was replaced with a 0.05% Triton X-100 solution for 3 hours. The tissues were then washed for 4 days by stirring in dH_2_O to remove any residual detergent. Decellularized tissue was sectioned and H&E stained, along with Western Blot to assess decellularization. Decellularized ECM or intact tissues were lysed in 25mM Tris, 150mM NaCl, 10% glycerol, 1% NP 40 and 0.5M EDTA with 1x protease Mini-complete protease inhibitor (04693124001; Roche, Indianapolis, IN) and 1x phosphatase inhibitor cocktail (4906845001; Roche, Indianapolis, IN) at 4°C. In addition, for decellularized ECM, tissue homogenization was performed using a BeadBug^TM^ 3 Place Microtube Homogenizer (D1030; Benchmark Scientific, Sayreville, NJ). Decellularized ECM or cell homogenate was then spun down at 21,000xg for 10min at 4°C and supernatant stored at −20°C until used for Western Blot, Coomassie or proteomics analysis.

### Sample preparation for mass spectrometry

Decellularized ECM homogenate (from ±5mm piece of tissue) was denatured in 8M urea and 10mM dithiothreitol and incubating on a shaker at 37°C for 2 hours. The samples were then alkylated with 25mM iodoacetamide and incubating at room temperature in the dark for 30 min. Following dilution to 2M urea in 11mM ammonium bicarbonate pH 8.0, deglycosylation was performed by adding 1000 units of PNGaseF (P0704S; New England Biolabs, Ipswich, MA), where after the samples were incubated on a shaker at 37°C for 2 hours. Samples were digested first using 1μg of endoproteinase LysC (125-05061; Wako Chemicals USA, Richmond, VA), shaking at 37°C for 2 hours, then adding 3μg trypsin (PR-V5113; Promega, Madison, WI) and shaking at 37°C overnight. The next day, additional 1.5μg trypsin was added to each sample and incubated on a shaker at 37°C for a further 2 hours. Samples were acidified with 50% trifluoroacetic acid and pH tested to <2.0 pH. Samples were centrifuged and supernatant collected. Peptide labeling with TMT 10plex (90110; Thermo Fisher Scientific, Waltham, MA) was performed per manufacturer’s instructions. Briefly, lyophilized samples were dissolved in ethanol and triethylammonium bicarbonate (TEAB), pH 8.5, and mixed with the TMT reagent. The resulting solution was vortexed and incubated at room temperature for 1 h. Samples labeled with the ten different isotopic TMT reagents were combined and concentrated to completion in a vacuum centrifuge. The peptides were fractioned via high-pH reverse phase HPLC. Peptides were resuspended in 100uL buffer A (10mM TEAB, pH8) and separated on a 4.6mm × 250 mm 300Extend-C18, 5um column (Agilent, Santa Clara, CA) using a 90min gradient with buffer B (90% acetonitrile (ACN), 10mM TEAB, pH8) at a flow rate of 1ml/min. The gradient was as follows: 1-5% B (0-10min), 5-35% B (10-70min), 35-70% B (70-80min), 70% B (80-90min). Fractions were collected over 75 minutes at 1min intervals from 10 min to 85 min. The fractions were concatenated into 15 fractions non-contiguously (1+16+31+46+61, 2+17+32+47+62, etc). The fractions were concentrated in a vacuum centrifuge and then were lyophilized. Lyophilized peptides were re-suspended in 100 uL of 0.1% formic acid and 10 uL was injected on to the LC-MS.

### LC-MS/MS analysis

Peptides were separated by reverse phase HPLC (Thermo Easy nLC1000) (Thermo Fisher Scientific, Waltham, MA) using a precolumn (made in house, 6 cm of 10 µm C18) and a self-pack 5 µm tip analytical column (12 cm of 5 µm C18; New Objective, Woburn, MA) over a 140 minute gradient before nanoelectrospray using a QExactive HF-X mass spectrometer (Thermo Fisher Scientific, Waltham, MA). Solvent A was 0.1% formic acid and solvent B was 80% ACN/0.1% formic acid. The gradient conditions were 2-10% B (0-3 min), 10-30% B (3-107 min), 30-40% B (107-121 min), 40-60% B (121-126 min), 60-100% B (126-127 min), 100% B (127-137 min), 100-0% B (137-138 min), 0% B (138-140 min), and the mass spectrometer was operated in a data-dependent mode. The parameters for the full scan MS were: resolution of 60,000 across 350-2000 m/z, automatic gain control 3e6, and maximum ion injection time of 50 ms. The full MS scan was followed by MS/MS for the top 15 precursor ions in each cycle with a normalized collision energy of 34 and dynamic exclusion of 30 s.

### Proteomics data processing and visualization

Raw mass spectral data files (.raw) were searched using Proteome Discoverer (Thermo Fisher Scientific, Waltham, MA) and Mascot version 2.4.1 (Matrix Science, Boston, MA). Mascot search parameters were: 10 ppm mass tolerance for precursor ions; 15 mmu for fragment ion mass tolerance; 2 missed cleavages of trypsin; fixed modification was carbamidomethylation of cysteine and TMT 10-plex modification of lysine and peptide N-Termini; variable modifications were methionine, lysine and proline oxidation, asparagine and glutamine deamination, Gln>pyro-flu(N-Term Q), carbamylation of N-terminus, tyrosine, serine and threonine phosphorylation. Only peptides with a Mascot score greater than or equal to 25 and an isolation interference less than or equal to 30 were included in the data analysis. Reporter ion intensities were median-centered and log2-transformed. Data was filtered for ECM protein groups with > 1 unique peptide and the resulting groups imported into Perseus data visualization and statistics software v1.6.1.1 (Tynova et al, 2016). Data visualization included scatterplot generation with coefficient of determination (R^2^) values to assess reproducibility between biological replicates and principal component analysis to assess differences between groups.

### Western Blot and Coomassie

Standard procedures were used for protein electrophoresis and western blotting. Protein lysates were separated by SDS-PAGE, transferred to a nitrocellulose membrane, blocked with 5% non-fat dry milk solution and incubated in primary antibody overnight at 4°C. Proteins were detected using HRP-conjugated secondary antibodies. For Coomassie, following SDS-PAGE separation, the gel was stained using Coomassie blue stain (20279; Thermo Fisher Scientific, Waltham, MA) solution, then destained overnight and imaged. Imaging was performed using a ChemiDoc^TM^ MP imaging system (12003154; Bio-Rad, Hercules, CA).

### Immunohistochemistry

Fixation, processing and staining of tissue sections from tumors and mammary fat pads was carried out as previously described^21^. Mammary fat pads dissected from C57BL/6 mice and tumors dissected from NOD-SCID mice were fixed in 10% buffered formalin and embedded in paraffin. For H&E staining: standard procedures were followed for H&E, including deparaffinizing, hydration, staining with Hematoxylin (GHS280; SIGMA, St. Louis, MO) and counterstaining with Eosin (HT110180; SIGMA, St. Louis, MO). For immunofluorescence: Tissue sections (10μm thick) were deparaffinized followed by antigen retrieval using Citra Plus solution (HK057; Biogenex, Fremont, CA). After blocking in PBS-0.5% Tween-20 and 10% serum, sections were incubated with primary antibodies overnight at 4°C and fluorescently labeled secondary antibodies at room temperature for 2 hours. DAPI (D1306; Thermo Fisher Scientific, Waltham, MA) was used to stain cell nuclei, and fluorochromes on secondary antibodies included AlexaFlour 488, AlexaFluor 534, AlexaFluor 647 (Jackson Immunoresearch, West Grove, PA). Sections were mounted in Fluoromount mounting media (00-4958-02; Thermo Fisher Scientific, Waltham, MA) and imaged using a Keyence BZ-X710 microscope (Keyence, Elmwood park, NJ).

### Adhesion assay

Depending on the experimental condition, cells were suspended in medium containing control antibody, vehicle (DMSO), inhibitory antibody or drug and plated on glass-bottomed dishes (MatTek, Ashland, MA) coated with 20μg/ml ECM protein and allowed to adhere for 2 hours. Cells were then fixed for 15 min in 4% paraformaldehyde, then permeabilized with 0.2% TritonX-100, blocked with 3% BSA and incubated with primary antibodies overnight at 4°C. Cells were DAPI (D1306; Thermo Fisher Scientific, Waltham, MA) and Phalloidin (A12390; Thermo Fisher Scientific, Waltham, MA) stained along with incubation with fluorescently labeled secondary antibodies at room temperature for 2 hours. Imaging was performed using a Keyence BZ-X710 microscope (Keyence, Elmwood park, NJ) and CellProfiler v3.1.8 (Carpenter et al) was used for imaging analysis using a custom pipeline. Image J (National Institutes of Health, Bethesda, MD) was used for pFAK foci counting. Data are the result of 3 independent experiments with 3 technical replicates per experiment.

### 2D migration assay and reseeding in decellularized ECM

For 2D migration, cells were plated on glass-bottomed dishes (MatTek, Ashland, MA) coated with 20μg/ml ECM protein and allowed to adhere for 1 hour. For reseeding on decellularized ECM, 5mm diameter pieces of decellularized tissue were cut and cells seeded and allowed to adhere for 6 hours. Depending on the experimental condition, media was then replaced with medium containing control antibody, vehicle (DMSO), inhibitory antibody, drug, growth factor or peptide. Cells were imaged overnight with images acquired every 10 min (for 2D migration assay) or 20 min (for reseeding in decellularized ECM) for 16 hours in an environmentally controlled chamber within the Keyence BZ-X710 microscope (Keyence, Elmwood park, NJ). Cells were then tracked using VW-9000 Video Editing/Analysis Software (Keyence, Elmwood park, NJ) and both cell speed and distance migrated calculated using a custom MATLAB script vR2018a (MathWorks, Natick, MA). Data are the result of 3 independent experiments with 6 fields of view per experiment and an average of 6 cells tracked per field of view (at least 100 cells per condition).

### Spheroid invasion assay

Cells were seeded in low-attachment plates in media, followed by centrifugation to form spheroids. Spheroids were grown for 3 days after which matrix was added to each well, which included (depending on the condition) Collagen I protein, Collagen VI protein, ETP, 10mM NaOH, 7.5% 10x DMEM and 50% 1x DMEM. The spheroids in matrix were then spun down and a further 50μl of media added to each well. Following another 4 days of growth, spheroids were imaged as a Z-stack using a Keyence BZ-X710 microscope (Keyence, Elmwood park, NJ) and Z-projection images analyzed using a Hybrid Cell Count feature within the BZ-X Analyzer software v1.3.1.1 (Keyence, Elmwood park, NJ). Data are the result of 3 independent experiments with 6 technical replicates per experiment.

### Cell viability assay

Cells were seeded in plates coated with 20μg/ml ECM protein and allowed to adhere for 6 hours. Medium was then changed to medium containing control antibody, vehicle (DMSO), inhibitory antibody or drug and incubated for 16 hours. PrestoBlue^TM^ Cell Viability Reagent (A13261; Invitrogen, Carlsbad, CA) was added to each well according to the manufacturer’s instructions and incubated for 25 min at 37°C. Fluorescence was then read on a plate reader at 562nm. Background was corrected to control wells containing only cell culture media (no cells). Data are the result of 3 independent experiments with 3 technical replicates per experiment.

### Statistical analysis

GraphPad Prism v7.04 was used for generation of graphs and statistical analysis. For data with normal distribution: for comparison between two groups, an unpaired two-tailed Student’s t-test was used and a p-value of ≤ 0.05 considered significant, and for comparison between multiple groups a one-way ANOVA with Bonferroni multiple testing correction was used with a corrected p-value of ≤ 0.0322 considered significant. For data with non-normal distribution: for comparison between two groups, an unpaired two-tailed Mann-Whitney test was used and a p-value of ≤ 0.05 considered significant, and for comparison between multiple groups a Kruskal-Wallis test with Dunn’s multiple testing correction was used with a corrected p-value of ≤ 0.0322 considered significant. Data represent mean ± SEM.

## Results

### ECM isolated from an obese mammary gland drives tumor cell invasion

We set out to understand whether obesity-driven changes in ECM could contribute to breast cancer invasion and how these effects compared to tumor ECM. We used a diet-induced model of obesity and two breast cancer models, xenografts generated with human triple-negative breast cancer (TNBC) cells in immunocompromised mice and the metastatic PyMT-MMTV genetic mouse model (Fig 1A). We isolated the mammary tissues of female C57Bl/6 mice fed on chow or 60%kcal high fat diet (HFD) for 16 weeks. Mice fed a HFD diet weighed twice as much as those fed with a chow diet and had 2-5g of subcutaneous fat (Fig S1A,B). H&E staining of mammary gland tissue sections confirm adipocyte hypertrophy in the tissue obtained from obese mice (Fig S1C). To investigate ECM-specific effects on tumor cell migration, we used previously established tissue decellularization methods to remove all cells within a tissue, while maintaining an intact ECM. Decellularization with low dose SDS removed all the nuclei, as verified by H&E staining (Fig 1B) and removed soluble intracellular proteins and enriched for ECM proteins (Fig S1D).

**Figure 1:**
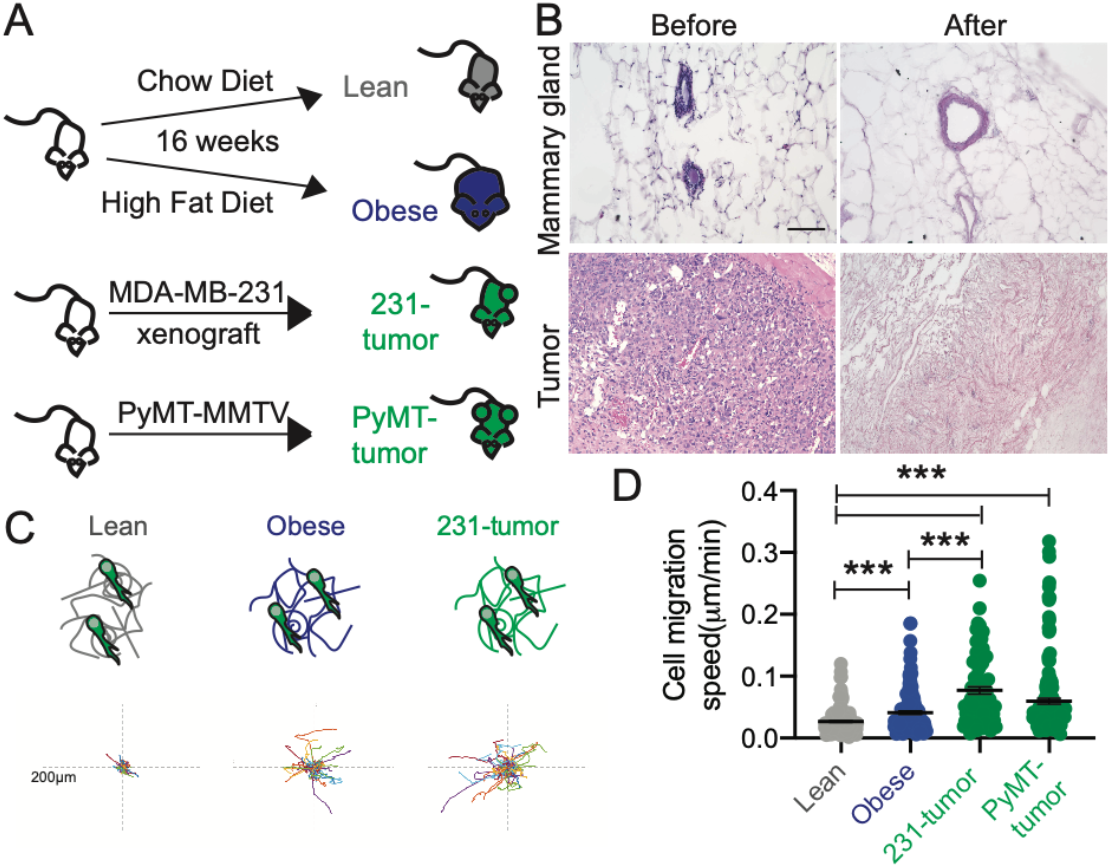
ECM isolated from the mammary gland from obese and tumor-bearing mice drives breast cancer cell invasion. A) Schematic of experimental setup: C57BL/6 mice were fed chow or high fat diet for 16 weeks. Two tumor models were used: MDA-MB-231 mammary xenografts grown in NOD-SCID mice for 8 weeks and tumors from the spontaneous PyMT-MMTV grown for 10 weeks. B) H&E sections of mammary gland and tumor before and after decellularization (Scale bar = 200µm). C) Schematic of experimental setup where GFP-labelled breast cancer cells were seeded on the decellularized ECM scaffolds isolated from the various mouse mammary glands with representative rose pots of cell migration. Each colored line represents the tracked movement of a cell over 16hrs. D) Cell migration speed of MDA-MB-231 cells stably expressing GFP (231-GFP) seeded on decellularized ECM obtained from lean mammary gland, obese mammary gland and 231-tumor xenograft and PyMT-GFP mouse cells derived from the spontaneous PyMT-MMTV tumors seeded onto decellularized ECM from PyMT-MMTV tumors. Each point represents the average speed of a cell over 16hrs. Data show mean ± SEM. A nonparametric Kruskal-Wallis test with Dunn’s multiple testing correction was performed, with *** p ≤ 0.005. Data obtained from at least 3 different ECM scaffolds from 3 different mice, with an average of 108 total cells per condition.

We seeded a GFP-labelled human TNBC cell line MDA-MB-231 on the ECM scaffolds from lean, obese and xenograft tumors, imaged them overnight and analyzed cell movement (Fig 1C). Decellularized ECM isolated from obese mammary glands significantly increased cell invasion speed relative to ECM from lean mammary glands (Fig 1D). As expected, tumor ECM also drives significant MDA-MB-231 cell invasion. Tumor ECM, obtained by the same decellularization method, also had a significant effect on cell speed relative to obese ECM (Fig 1C,D). We seeded GFP-labelled PyMT tumor-derived cells in the ECM isolated from primary MMTV-PyMT tumors, which also increased tumor cell invasion (Fig 1D). These data demonstrate that obese ECM drives tumor cell invasion and that there may be ECM proteins commonly upregulated in both tumor and obese ECM that may drive similar effects on tumor cell migration.

### Obesity is associated with changes in ECM composition of mammary gland tissue

To comprehensively characterize the changes in ECM associated with obesity, we performed proteomics analysis. To directly compare the abundance of each peptide within the different samples, we used label-based quantification with unique isobaric tandem mass tags (TMT) to label 3 samples per group. We used the existing in silico murine matrisome data to annotate the data and identify proteins with differential abundance in obese vs. lean mammary tissues. We first assessed reproducibility between biological replicates by plotting median-centered reporter ion intensities and found significant correlation between all combinations of individual samples within each group (Fig S2). In the matrisome of the obese mammary gland, we identified 53 core matrisome proteins (27 glycoproteins, 18 collagens and 8 proteoglycans) and 15 ECM-associated proteins (10 ECM-affiliated proteins, 3 ECM regulators and 2 secreted factors) (Fig S3). These studies provide the first characterization of the matrisome of obese mammary tissues. They demonstrate that the ECM composition of lean and obese mammary tissues is distinct (Fig 2A,B). We then compared the obese matrisome to two previously published mammary gland tumor proteomics^18,19^ (Fig 2C). We identify a common signature of 9 matrisome proteins, which are upregulated in both tumor and obese ECM, which includes Collagen VI and XXII and FN, LAMA5, VTN, ELN, VWA1, LGALS1 and ANXA3 (Fig 2D).

**Figure 2:**
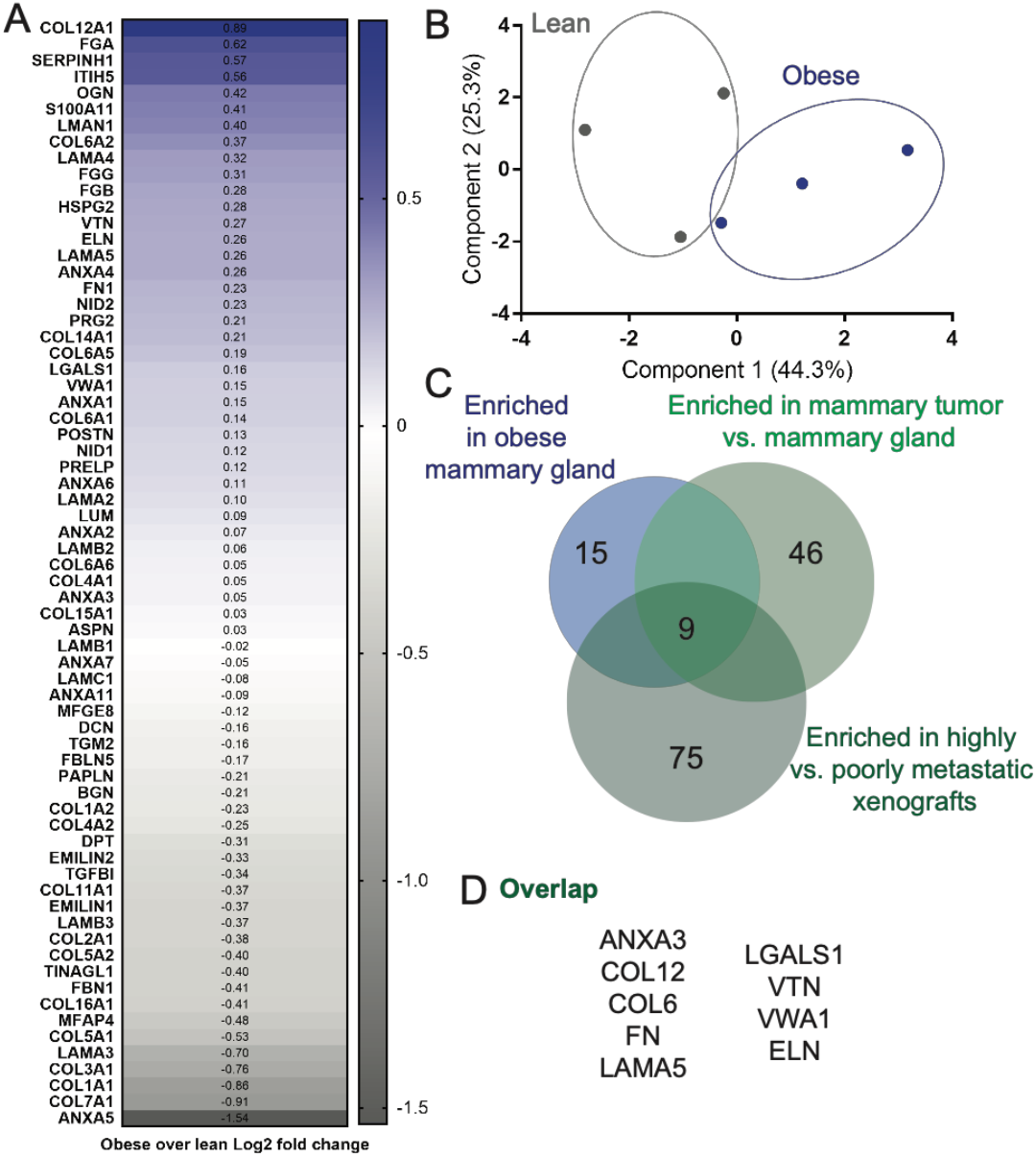
Proteomics analysis of ECM isolated from obese mammary gland reveals overlap with ECM from mammary tumors. Obese and lean mammary gland ECM were compared through average log2 fold change differences of 68 ECM proteins (A) and principal components analysis (PCA) (B), showing a number of enriched proteins in obese mammary gland ECM and the clustering of obese and lean samples. C) ECM proteins found enriched in obese mammary gland were overlapped with proteins enriched in 4T1 orthotopic syngeneic mammary tumor (1) and in MDA-MB-231-LM2 highly metastatic xenografts (2) in a Venn diagram. D) List of 9 matrisome proteins present in obese mammary gland ECM, 4T1 tumor ECM and metastatic LM2 tumors.

### Collagen VI, which is upregulated in obese and tumor ECM, drives migration of TNBC cancer cells

Our next goal was to identify which of the ECM proteins upregulated both in tumor and obese ECM contributes to tumor cell invasion. From the 9 ECM proteins identified above, we chose to focus on 4: Fibronectin, Collagen VI, Elastin and Laminin, each of these proteins is commercially available and has published antibodies for staining. We used 3 main assays commonly used to study cell-ECM interactions: an adhesion assay, 2D cell migration and 3D invasion. For adhesion assays, MDA-MB-231 cells were seeded on the 4 ECM proteins and left to adhere for 2hrs (Fig 3A). Collagen VI increased both cell area and cell eccentricity (Fig S4A,B). FN increased cell area, but did not alter cell shape. Neither Laminin nor Elastin had any significant effect on cell shape. Similar results were obtained with another TNBC cell line MDA-MB-468 (Fig S4C-E). We then plated the cells overnight on the 4 ECM proteins to evaluate their effects on cell migration speed. Here, we find that Collagen VI significantly increases cell migration of both MDA-MB-231 and MDA-MB-468 cells (Fig 3C,D). To investigate whether these effects on 2D cell migration are tumor cells specific, we also performed the same assay on the epithelial breast cancer cell line MCF10A. We find that Collagen VI also increases cell migration of MCF10A cells (Fig S4F). We confirmed that Collagen VI could induce cell migration of breast cancer cells at a range of concentrations (Fig S4G). We then investigated whether Collagen VI could also drive 3D invasion of breast cancer cells. We generated spheroids embedded in Collagen I, the most abundant ECM protein present in normal breast and breast tumor tissue. We find that Collagen VI significantly increases MDA-MB-231 invasion in a 3D spheroid model (Fig 3E,F). Overall, these data identify Collagen VI, an ECM protein upregulated in tumor and obese ECM, as a major driver of breast cancer cell migration.

**Figure 3:**
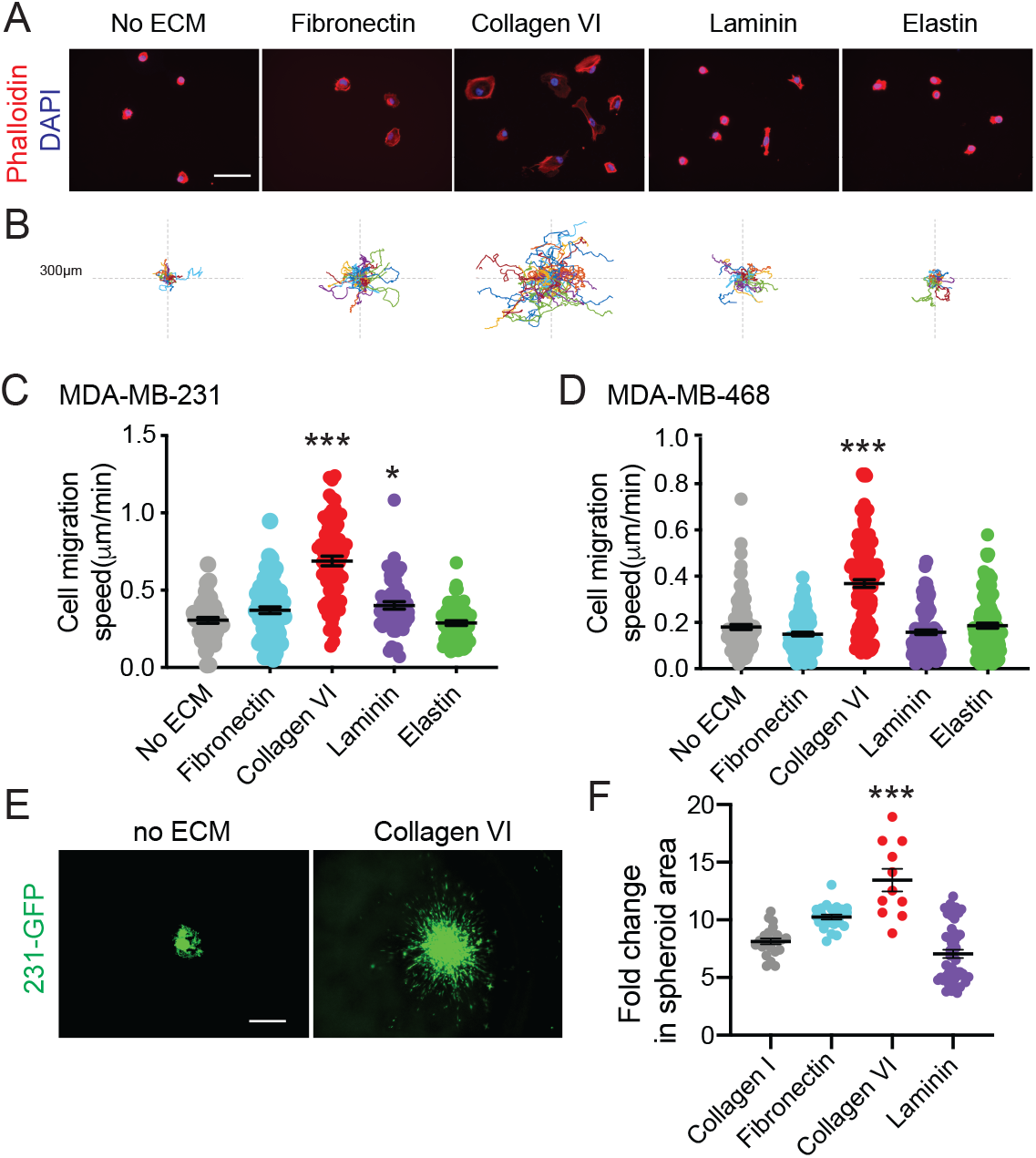
Collagen VI, ECM protein upregulated in obese and tumor ECM, drives breast cancer cell adhesion, migration and invasion. A) Representative images from an adhesion assay of MDA-MB-231 cells on 20µg/ml Fibronectin, Collagen VI, Laminin or Elastin (Scale bar = 50µm). B) Representative rose plots of cell migration tracks of MDA-MB-231 cells on ECM substrates, with each colored line representing a single cell track over 16h time course. Speed of MDA-MB-231 (C) and MDA-MB-468 (D) cells seeded on 20µg/ml Fibronectin, Collagen VI, Laminin or Elastin. Each point represents the average speed of a cell over 16hrs. E) Representative images of spheroid 231-GFP (MDA-MB-231 GFP-labelled cells) cells after 5 days within media without ECM and in a Collagen I + Collagen VI (100µg/ml) matrix. (Scale bar = 300µm). F) Quantification of fold change in 231-GFP spheroid area after 5 days. Data show individual cells or spheroids and mean ± SEM. Significance was determined by a nonparametric Kruskal-Wallis test with Dunn’s multiple testing correction was performed, with *p<0.05, **p<0.01 and ***p<0.005. For each experiment, data pooled from at least 3 independent experiments.

### Collagen VI is upregulated in obese and tumor ECM

Collagen VI^20^ is secreted by stromal adipocytes in breast tissue^22^. We performed immunostaining for Collagen VI in the lean and obese mammary glands obtained from mice descried in Figure 1. We found increased accumulation of Collagen VI in obese mammary gland relative to lean (Fig 4A,B). Collagen VI is encoded by six different genes (COL6A1 to A6) that each code for α chain. Collagen VI assembles into a triple-helix monomer composed of α1–α2–αX chains, where αX is either α3,4,5 or 6^23,24^. We mined the TCGA dataset^25^ and found that patients with high levels of α1–α2–α3 chains have worse outcomes than those with low levels (Fig 4C). Finally, we confirmed expression of Collagen VI in human triple-negative breast cancer, with high levels of Collagen VI in the stroma of human tumors (Fig 4D). These experiments confirm that Collagen VI in obese ECM and is associated with poor patient outcomes.

**Figure 4:**
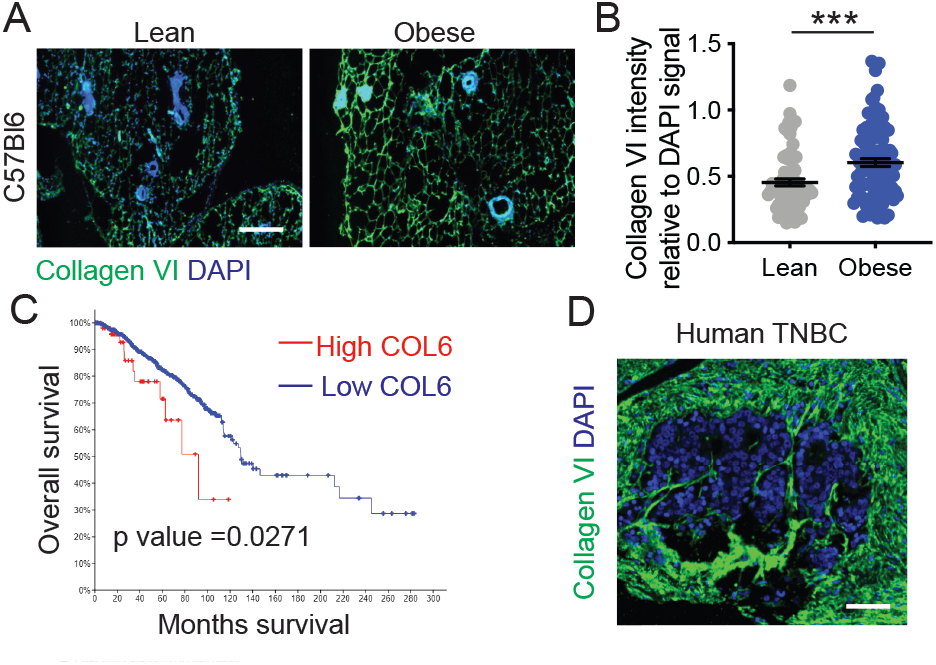
Collagen VI is enriched in obese mammary gland ECM and patients with high tumor Collagen VI have lower overall survival. A) Immunostaining of mammary gland sections from lean and obese C57BL/6 mice stained for Collagen VI (green) and nuclei (blue) (Scale bar = 100µm). B) Collagen VI intensity relative to DAPI signal shows increased abundance in obese ECM. Unpaired two-tailed Mann-Whitney test performed. Data represent mean ± SEM. *** p ≤ 0.005. C) Kaplan-Meier curve of human breast cancer patients with low vs high Collagen VI (COL6), data from TCGA. D) Collagen VI (green) and DAPI (Blue) staining of a human TNBC tissue array indicates the presence of Collagen VI in stroma of human tumors (Scale bar = 75µm).

### Collagen VI drives migration of TNBC cancer cells via NG2 and EGFR crosstalk and MAPK signaling

In breast cancer, Collagen VI has been suggested to drive breast cancer cell migration via its C-terminal fragment, endotrophin (ETP). Endotrophin (ETP) is a cleavage product of the COL6α3 chain^24^ (Fig 5A). Previous studies have found that ETP is abundant in tumor tissues in PyMT mice, and that ETP enhances fibrosis, angiogenesis, inflammation, and the epithelial to mesenchymal transition (EMT), thereby promoting primary tumor growth and metastasis. ETP is thought to serve as the major mediator of the Collagen VI-mediated tumor effects in breast cancer^22^. We compared effects of Collagen VI and ETP in our assays, both by embedding ETP in the matrix and including it as a soluble factor in the solution, alone or in combination with the Collagen I. ETP did not have an effect on the cell migration of MDA-MB-231 cells (Fig 5B,C). Furthermore, we find that the ETP fragment did not induce MDA-MB-231 cell migration, either attached, bound to Collagen I, or soluble (Fig 5C). In other contexts, Collagen VI can signal through the proteoglycan NG2, β-1 integrins as well as via crosstalk with RTKs (Fig 5A)^24^. We find that Collagen VI-driven MDA-MB-231 breast cancer cell migration is partially inhibited by an NG2 specific antibody (NG2ab), but not by β1 inhibitory antibody (β1ab) (Fig 5D). Neither antibody affected proliferation at the doses used (Fig S5A). We confirmed these findings in MDA-MB-468 cells (Fig S5B). Because NG2 binds to the central portion of Collagen VI and not the c-terminal ETP fragment, these data demonstrate for the first time that Collagen VI drives breast cancer invasion through an ETP-independent mechanism. NG2 has been shown to crosstalk with RTKs such as EGFR^26^ to drive intracellular signaling. Treatment with NG2ab and the EGFR inhibitor Lapatinib, individually or in combination, reduced cell adhesion to Collagen VI relative to their own vehicle controls (Fig 5E), at a dose that did not affect cell viability (Fig S5C). Treatment with Lapatinib also inhibited Collagen VI-driven migration, alone and in combination with NG2ab (Fig 5F). Together, these studies suggest that NG2 and EGFR synergize to mediate Collagen VI-driven breast cancer cell adhesion and migration.

**Figure 5:**
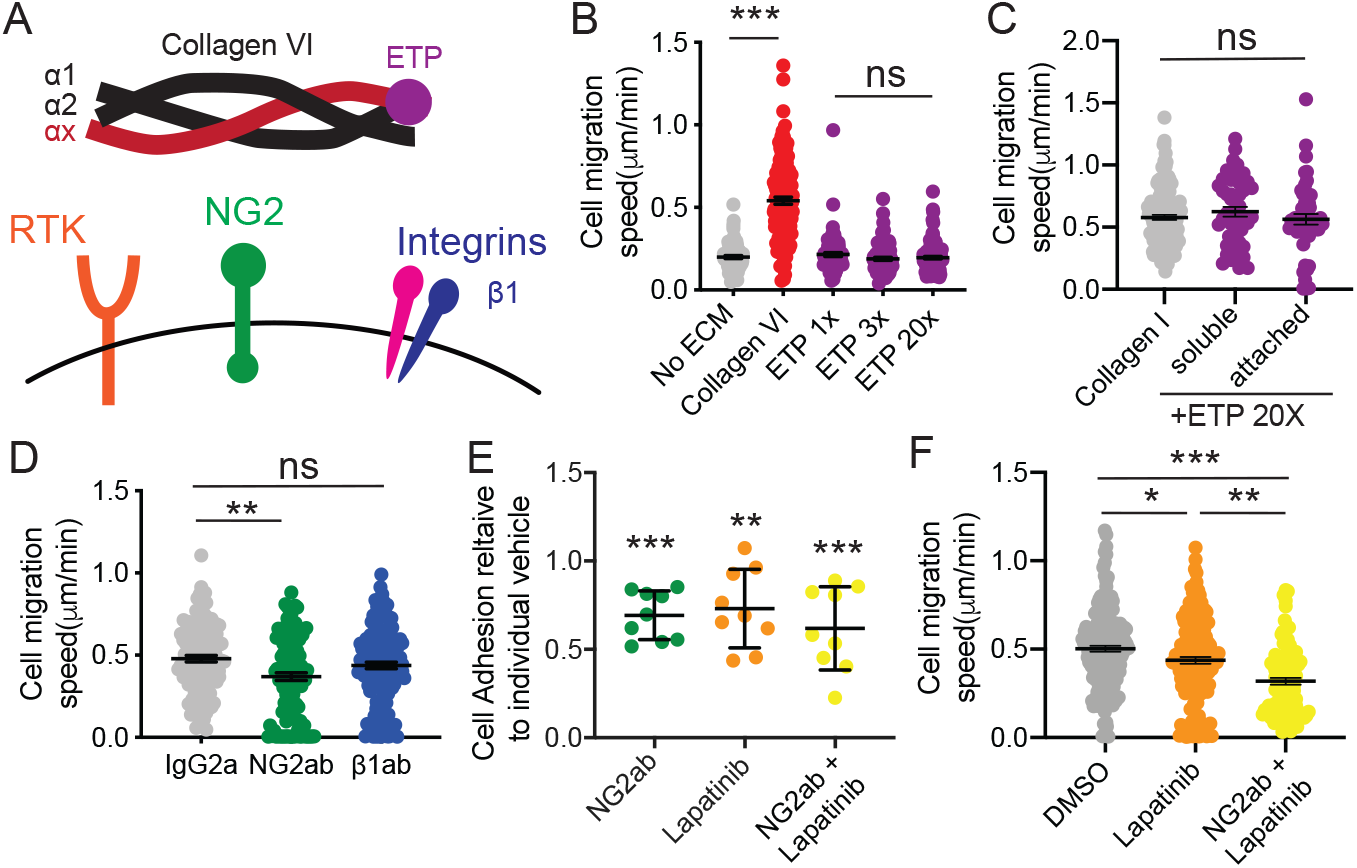
Collagen VI-driven breast cancer migration is mediated by NG2 and EGFR crosstalk. A) Schematic of structure of Collagen VI, which assembles into a triple-helix monomer composed of α1–α2–αX chains, where αX is either α3,4,5 or 6. Collagen VI can bind to NG2, β1 integrins bind and NG2 can co-signal with EGFR. Endotrophin (ETP) is a small terminal peptide on the α3 chain. B) Cell migration speed of MDA-MB-231 cells on full length Collagen VI or ETP (at 1x, 3x and 20x the relative amount to the ETP present on a full length Collagen VI strand) C) Cell migration speed of MDA-MB-231 cells seeded on Collagen I in combination with ETP or with ETP in the media. D) Cell migration speed of MDA-MB-231 cells seeded on 20µg/ml Collagen VI and treated with IgG2a, NG2 targeting antibody (NG2ab) or β1 integrin inhibitory antibody (β1ab) E) MDA-MB-231 cell adhesion in the presence of NG2ab, EGFR inhibitor Lapatinib (10μM) or a combination of both. Data show cell adhesion relative to respective vehicle conditions: IgG2a, DMSO or the combination. F) Cell migration speed of MDA-MB-231 cells seeded on 20µg/ml Collagen VI and treated with DMSO, Lapatinib (10μM) and combination of both. Graphs in B-D and F show individual cell data and mean ± SEM. Significance was determined by a nonparametric Kruskal-Wallis test with Dunn’s multiple testing correction, with *p<0.05, **p<0.01 and ***p<0.005. For each experiment, data pooled from at least 3 independent experiments.

Next, we investigated the downstream signaling pathways which drive Collagen VI-mediated responses. In neuronal tissues, Collagen VI acts via Akt, JNK, ERK and p38 MAPKs^27,28^. In sarcoma, Collagen VI activates PI3K signaling to drive cell migration^29^, whereas in breast cancer Collagen VI activates Akt and β-catenin to drive cell proliferation^20^. We investigated the effects of Alpelisib, a PI3K inhibitor, Defactinib, a FAK inhibitor and Trametinib, an MEK inhibitor, on cell migration at concentrations of 10 µM, 0.1 µM and 0.1 µM respectively. Cell viability assays were used to determine the appropriate dosage of the drugs investigated (Fig S5 D-F). Trametinib but, not Apelisib or Defactinib resulted in a significant decrease in Collagen VI-driven cell migration (Fig 6B), indicating that MAPK signaling downstream of EGFR and NG2 mediates Collagen VI-driven cell migration. Further, inhibition of EGFR and NG2 decreased phospho-p44/42 ERK signal in MDA-MB-231 cells seeded on Collagen VI (Fig 6C), without affecting pFAK397 (Fig S5G). These data demonstrate that Collagen VI drives migration in breast cancer cells via NG2/EGFR crosstalk and MAPK signaling.

**Figure 6:**
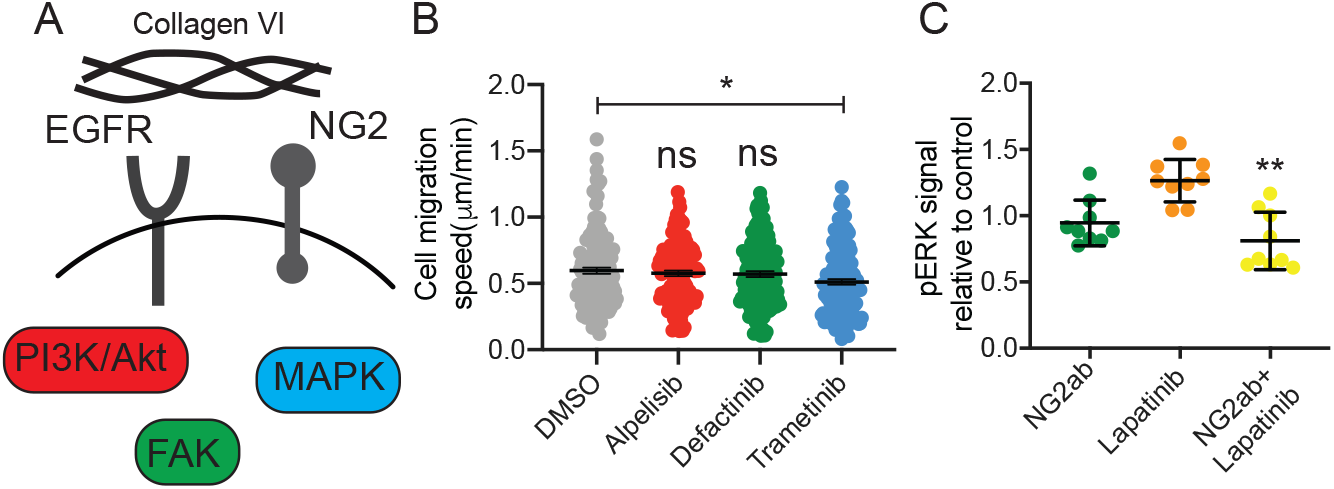
Collagen VI-driven migration is driven by MAPK signaling in breast cancer cells. A) Schematic of signaling pathways known to be activated downstream of NG2 and Collagen VI. B) Cell migration speed of MDA-MB-231 cells plated on Collagen VI and treated with Alpesilib (PI3K inhibitor, 10µM), Defactinib (FAK inhibitor, 0.1µM) and Trametinib (MEK inhibitor, 0.1µM). Graph shows individual cell data and mean ± SEM. Significance was determined by a nonparametric Kruskal-Wallis test with Dunn’s multiple testing correction, with *p<0.05. For each experiment, data pooled from at least 3 independent experiments. C) Immunostaining of phospho-p44/42 ERK (Thr202/Tyr204) in MDA-MB-231 cells seeded on Collagen VI (20µg/ml) and treated with NG2ab, Lapatinib and combination NG2ab and Lapatinib. Data represent mean ± SEM, statistics by one-way ANOVA *p<0.05, ** p ≤ 0.01.

### Blocking Collagen VI signaling reduces obese and tumor ECM driven invasion

Finally, we investigated whether Collagen VI-driven signaling was responsible for the increase in breast cancer cell invasion seen in decellularized ECM from tumor-bearing and obese mice (Fig 1). We seeded MDA-MB-231 cells treated with NG2ab in decellularized ECM from lean, obese and tumor-bearing mice. We found that inhibition of NG2 decreased migration in decellularized ECM obtained from both obese and tumor-bearing mice (Fig 7A,B). These studies demonstrate that Collagen VI-driven signaling contributes to ECM-mediated breast cancer cell invasion in the context of obesity and tumor progression.

**Figure 7:**
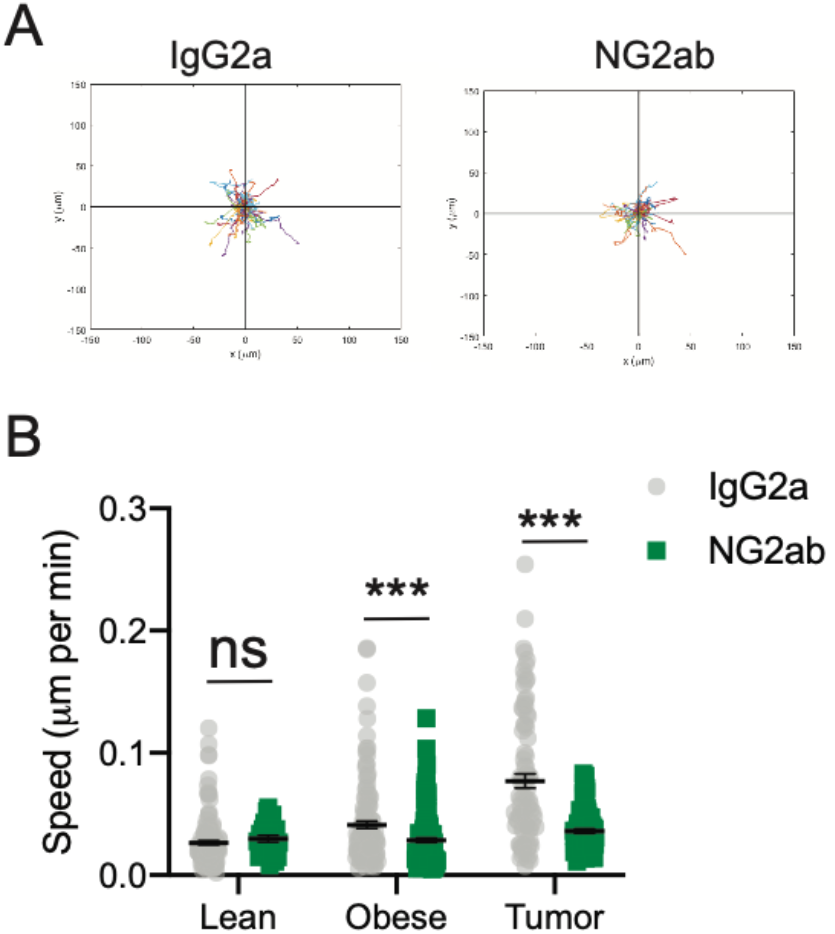
Inhibition of Collagen VI receptor NG2 reduces breast cancer cell migration on mammary gland ECM derived from obese mice and tumor ECM. A) Representative rose plots of 231-GFP cell migration seeded on decellularized ECM from obese mammary gland and treated with IgG2a or NG2ab, where each colored line represents the tracked movement of a cell over the time course (16hrs) B) Cell migration speed of 231-GFP cells seeded on mammary gland ECM derived from lean mice, obese mice and tumor-bearing mice treated with IgG2a or NG2ab. Graphs show mean ± SEM, with dot representing data from a single cell. Unpaired two-tailed Mann-Whitney tests were performed between respective IgG2a control and NG2ab conditions. Data obtained from at least 3 different ECM scaffolds from 3 different mice, with an average of 108 total cells per condition. Significance by t-test, with *** p<0.005

## Discussion

It is well established that obesity is associated with increased breast cancer incidence, metastasis and chemoresistance. Obesity is also known to be associated with fibrosis; how these changes in ECM properties impacts breast tumor progression remains poorly understood. Here, we characterize the ECM of the obese mouse mammary gland and find significant overlap with published mammary tumor ECM datasets. We identify Collagen VI as upregulated in both tumor and obese ECM and show that Collagen VI promotes breast cancer cell adhesion, 2D migration and 3D invasion via ECM and growth factor crosstalk and MAPK signaling. Overall, these studies define a novel mechanism by which obesity may contribute to breast cancer progression and identify Collagen VI as novel driver of breast cancer cell invasion.

Collagen VI is expressed in a range of tissues and has been shown to be involved in cartilage, bone and skeletal muscle function, in nervous system regeneration and myelination, and immune cell recruitment and polarization^24^. Previous studies have found that Collagen VI plays an important role in stimulating mammary tumor growth and breast cancer progression^20^. In MMTV-PyMT Collagen VI -/- mice, the lack of Collagen VI reduced hyperplasia and primary tumor growth. In this study, the authors state that Collagen VI has no effect on metastasis. However, the only method used to assess metastasis in this study was by injecting Met1 cells into the tail vein of wild-type or Collagen VI -/- mice and quantifying homing to the lungs. This assay evaluates whether Collagen VI is important for extravasation and metastatic outgrowth, but does not address the role of Collagen VI in driving local invasion in the primary tumor. Here, we demonstrate for the first time that Collagen VI directly drives cell migration of breast cancer cells. A subsequent study demonstrated that ETP, a cleavage product of the COL6α3 chain, serves as the major mediator of the Collagen VI-driven effects on cell migration and metastasis^22^, by enhancing TGFβ signaling to promote epithelial-mesenchymal transition (EMT), although the cell surface receptors and signaling pathways activated downstream of ETP are not known. Here, we do not see an effect of ETP on breast cancer cell migration, although this may be because MDA-MB-231 cells are mesenchymal. We find that Collagen VI-driven cell migration is partially inhibited by an NG2-specific antibody. Because NG2 binds to the central portion of Collagen VI and not the C-terminal ETP fragment, these data demonstrate for the first time that Collagen VI drives breast cancer invasion via NG2, and crosstalk with a receptor tyrosine kinase EGFR. Several ECM proteins are known to signal via adhesion receptor/RTK crosstalk, and it will be interesting to further dissect the mechanisms that mediate this and whether Collagen VI synergizes with growth factors to promote invasion in breast cancer.

While our results demonstrate how obesity-induced changes in ECM can contribute to tumor progression, we still do not know what cells secrete ECM in the adipose mammary gland. They also provide an experimental framework to study ECM-driven effects on tumor cells, in an unbiased manner, irrespective of the cell type which secreted it. Both the tumor and stroma compartment are known to contribute to ECM in mouse models of metastasis, via species-specific peptide identification^19^. Collagen VI is known to be secreted by adipocytes, and Collagen VI secretion increases during adipose differentiation^30^. However, adipocytes can secrete ECM proteins directly as well as can factors such as TGFβ which activates other cells such as myofibroblasts or macrophages to synthesize ECM proteins^31^. In the Seo *et al*., study, adipose stromal cells were isolated and found to have myofibroblast-like characteristics, as indicated by alpha-SMA staining^14^. It will be important to dissect which cell types are contributing to the changes in ECM associated with obesity in the mammary gland, which may reveal novel therapeutic strategies for targeting ECM-driven signaling that promotes tumor progression and metastasis. Finally, obese breast cancer patients have been shown to be more resistant to chemotherapy, the main standard of care of metastatic breast cancer. Obesity has been shown to contribute to chemoresistance by altering drug pharmacokinetics, inducing chronic inflammation, as well as altering tumor-associated adipocyte adipokine secretion ^32^. The role of obesity-driven ECM production in chemoresistance remains unknown. Interestingly, Collagen VI has been shown to drive resistance to chemotherapeutic drug cisplatin in ovarian cancer^33^. Future studies investigating the role of obesity-induced changes in ECM in drug resistance may also shed light on novel strategies to treat this growing population of breast cancer patients.

## Supplementary Figures

**Figure S1:**
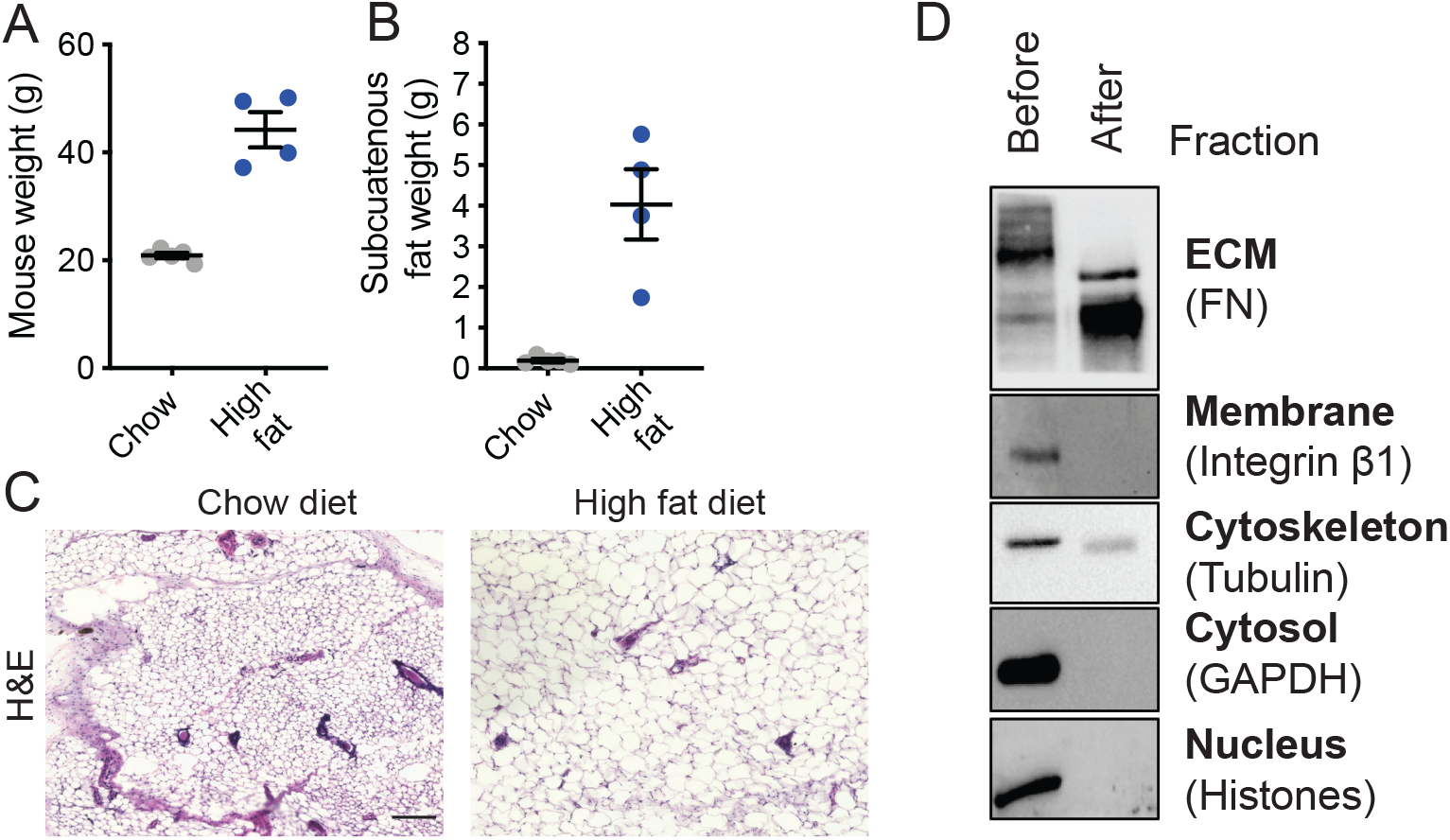
Assessment of diet-induced obesity and decellularization methods. C57BL/6 mice were fed on chow or high fat diet for 16 weeks after which both total body weight (A) and subcutaneous fat weight (B) were measured. C) H&E staining sections of mammary gland from chow and high fat diet fed mice were H&E processed, revealing significant hypertrophy of adipocytes (typical of fatty tissue) in the glands of mice fed with high fat diet (Scale bar, 200µm). D) Representative Western Blot of mammary gland tissue before and after the decellularization procedure. Enrichment in FN (representing ECM) and depletion of Integrin β1 (membrane), tubulin (cytoskeleton), GAPDH (cytosol) and histones (nucleus) reflects successful decellularization.

**Figure S2:**
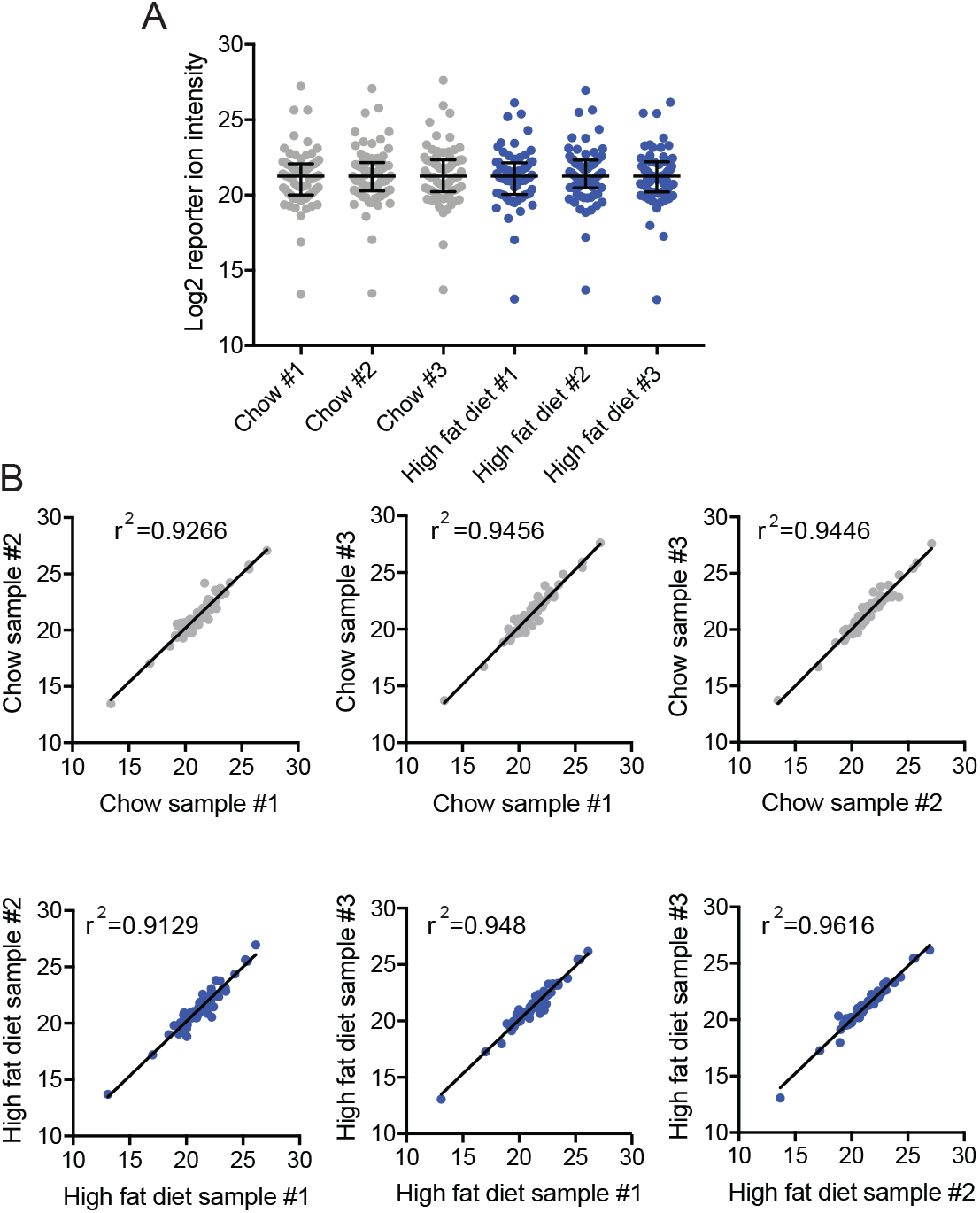
ECM proteomics data normalization and biological group correlations. **A)** ECM reporter ion intensities were median-centered and log2-transformed, the bars represent median with interquartile range. This normalizes the data, correcting for any potential inter-sample protein loading variances on the mass spectrometer, to enable accurate comparisons between abundance of ECM proteins. **B)** Scatterplots with r^2^ correlation values between biological replicates reveal a strong correlation between ECM proteins within the chow diet fed mice group and also between ECM proteins within the high fat diet fed mice group (values on x and y axes are log2 of reporter ion intensity).

**Figure S3:**
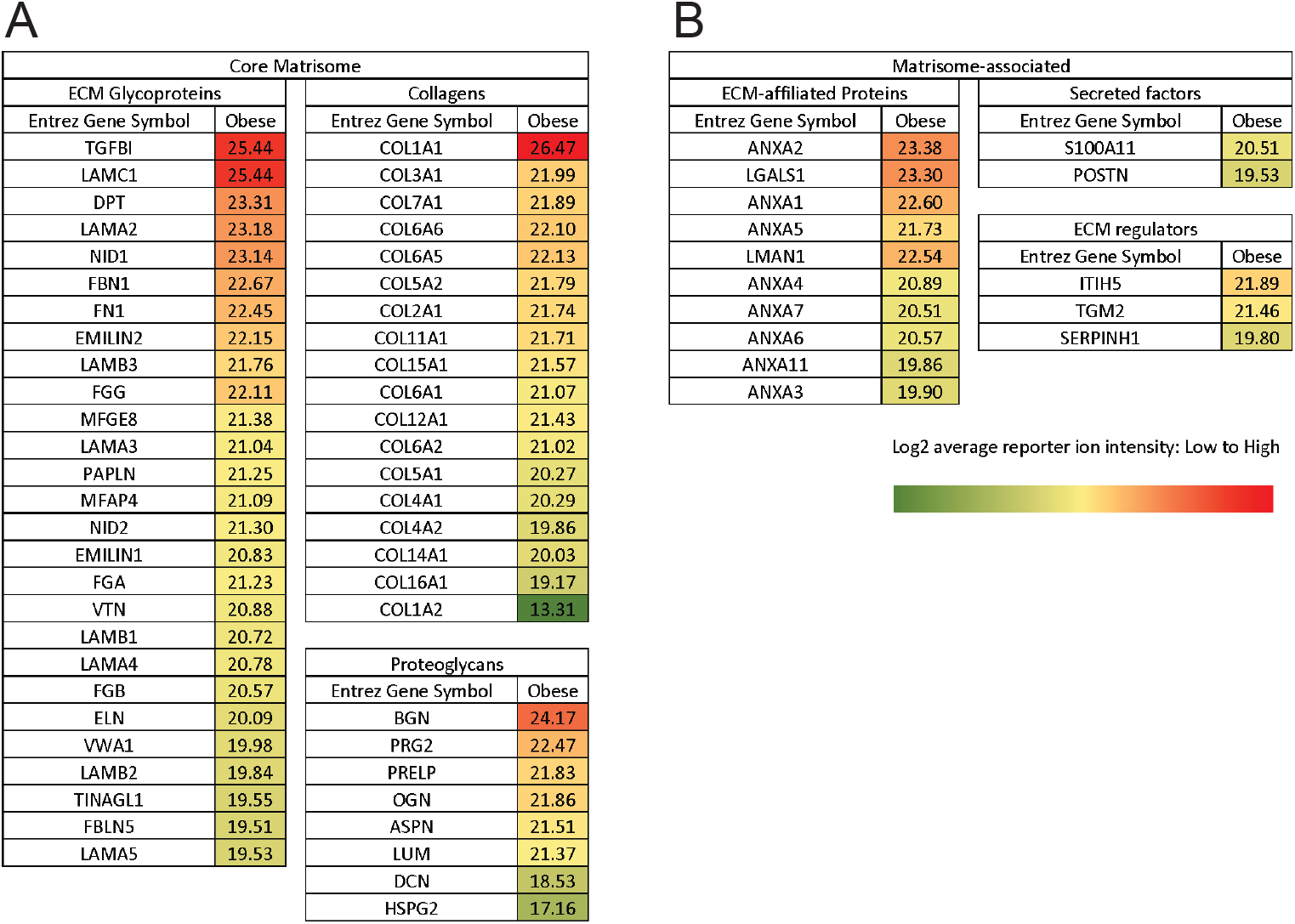
Matrisome profile of obese mammary fat pad. A) Core matrisome list, including ECM glycoproteins, collagens and proteoglycans. B) Matrisome-associated list including ECM-affiliated proteins, secreted factors and ECM regulators. Proteins were annotated according to the Matrisome Project. Data are the result of 3 biological replicates with the average log2 reporter ion intensity reported.

**Figure S4:**
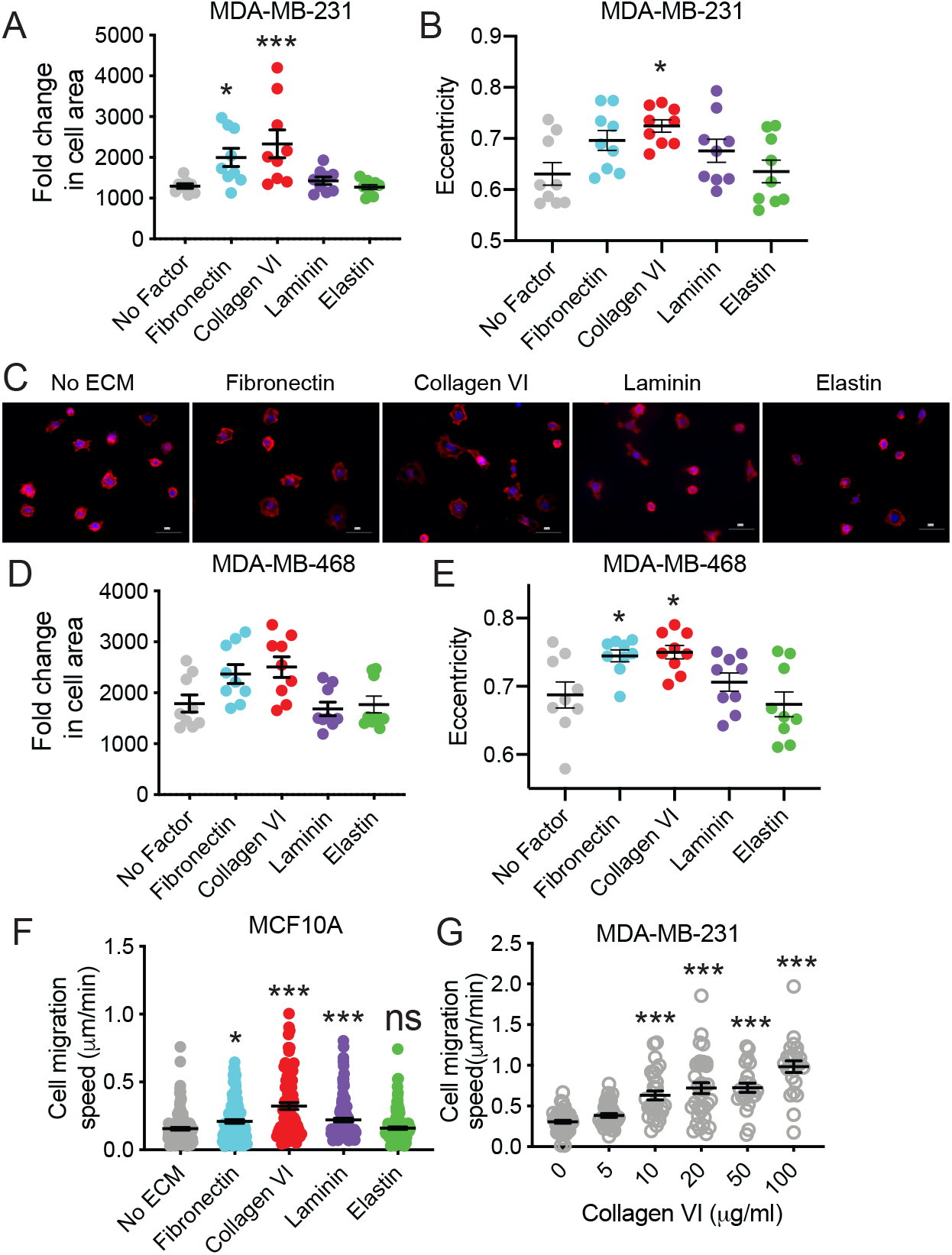
Collagen VI drives adhesion and migration of several breast cancer cell lines. Quantification of cell area (A) and cell eccentricity (B) for MDA-MB-231 cells on 20µg/ml Fibronectin, Collagen VI, Laminin or Elastin and allowed to adhere for 2hrs. C) Representative images of MDA-MB-468 breast cancer cells seeded on the different substrates (Scale bar = 50µm). Quantification of cell area (D) and cell eccentricity (E) for MDA-MB-468 cells on 20µg/ml Fibronectin, Collagen VI, Laminin or Elastin and allowed to adhere for 2hrs. Graphs show data from 3 independent experiments and average from 3 different fields of view per experiment. Significance was determined by a nonparametric Kruskal-Wallis test with Dunn’s multiple testing correction, with *p<0.05, ***p<0.005. F) Cell migration speed of pre-neoplastic mammary epithelial cell line MCF10A cells on 20µg/ml Fibronectin, Collagen VI, Laminin or Elastin and allowed to adhere for 2hrs. Each point represents the average speed of a cell over the time course (16hrs). G) Cell migration speed of MDA-MB-231 cells on different concentrations of Collagen VI. Graphs show individual cell data and mean ± SEM. Significance was determined by a nonparametric Kruskal-Wallis test with Dunn’s multiple testing correction, with *p<0.05 and ***p<0.005. For each experiment, data pooled from at least 3 independent experiments.

**Figure S5:**
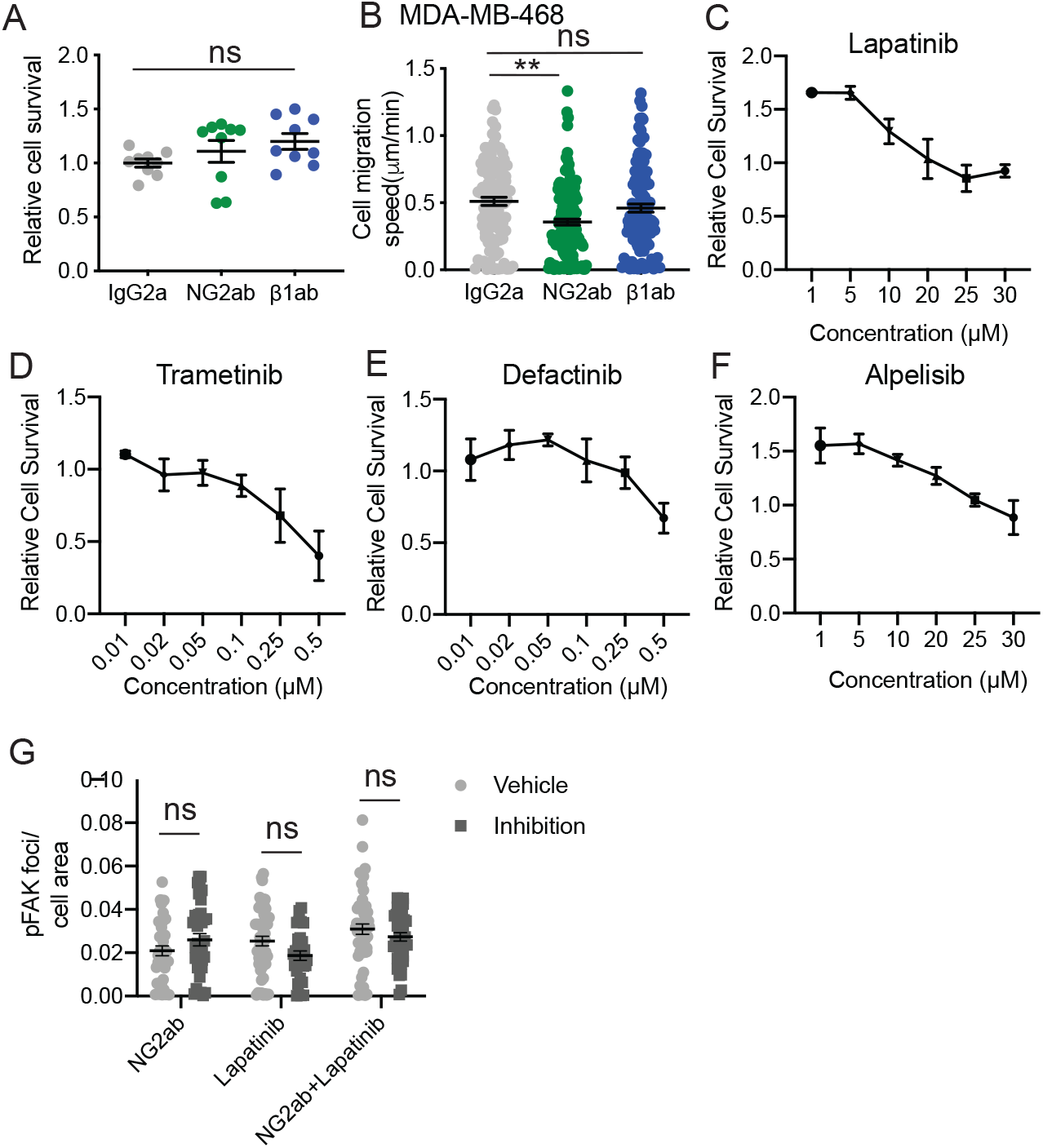
Investigation into receptors and signaling pathways mediating Collagen VI-driven cancer cell migration. A) Cell migration speed of MDA-MB-468 cells seeded on 20µg/ml Collagen VI and treated with IgG2a, NG2 targeting antibody (NG2ab) or β1 integrin inhibitory antibody (β1ab). Graph shows individual cell data and mean ± SEM. Significance was determined by a nonparametric Kruskal-Wallis test with Dunn’s multiple testing correction, with **p<0.01. For each experiment, data pooled from at least 3 independent experiments. B) Cell survival of MDA-MB-231 cells treated with IgG2a, NG2 targeting antibody (NG2ab) or β1 integrin inhibitory antibody (β1ab) for 16hrs. Dose response curves plotting the effect of C) Lapatinib D) Trametinib E) Defactinib and F) Apelisib on MDA-MB-231 cell survival after 16hrs. G) Number pFAK397 in breast cancer cells plated on Collagen VI for 2hrs and treated with NG2ab, EGFR inhibitor Lapatinib (10μM) or a combination of both. Data pooled for 3 experiments.

## References

1. Mitchell, S. & Shaw, D. The worldwide epidemic of female obesity. Best Pract. Res. Clin. Obstet. Gynaecol. (2015). doi:10.1016/j.bpobgyn.2014.10.002

2. Chen, S. et al. Obesity or overweight is associated with worse pathological response to neoadjuvant chemotherapy among Chinese women with breast cancer. PLoS One (2012). doi:10.1371/journal.pone.0041380

3. Bastarrachea, J., Hortobagyi, G. N., Smith, T. L., Kau, S. W. C. & Buzdar, A. U. Obesity as an adverse prognostic factor for patients receiving adjuvant chemotherapy for breast cancer. Ann. Intern. Med. (1994). doi:10.7326/0003-4819-120-1-199401010-00004

4. Ewertz, M. et al. Effect of obesity on prognosis after early-stage breast cancer. J. Clin. Oncol. (2011). doi:10.1200/JCO.2010.29.7614

5. Osman, M. A. & Hennessy, B. T. Obesity correlation with metastases development and response to first-line metastatic chemotherapy in breast cancer. Clin. Med. Insights Oncol. (2015). doi:10.4137/CMO.S32812

6. Sun, H. et al. Triple-negative breast cancer and its association with obesity. Mol. Clin. Oncol. (2017). doi:10.3892/mco.2017.1429

7. Fontanella, C. et al. Impact of body mass index on neoadjuvant treatment outcome: a pooled analysis of eight prospective neoadjuvant breast cancer trials. Breast Cancer Res. Treat. (2015). doi:10.1007/s10549-015-3287-5

8. Chen, X. et al. Obesity and weight change in relation to breast cancer survival. Breast Cancer Res. Treat. (2010). doi:10.1007/s10549-009-0708-3

9. Bianchini, G., Balko, J. M., Mayer, I. A., Sanders, M. E. & Gianni, L. Triple-negative breast cancer: Challenges and opportunities of a heterogeneous disease. Nature Reviews Clinical Oncology (2016). doi:10.1038/nrclinonc.2016.66

10. Anders, C. K., Zagar, T. M. & Carey, L. A. The management of early-stage and metastatic triple-negative breast cancer: a review. Hematol. Oncol. Clin. North Am. 27, 737–49–viii (2013).

11. Quail, D. F. & Dannenberg, A. J. The obese adipose tissue microenvironment in cancer development and progression. Nature Reviews Endocrinology (2018). doi:10.1038/s41574-018-0126-x

12. Hoy, A. J., Balaban, S. & Saunders, D. N. Adipocyte–Tumor Cell Metabolic Crosstalk in Breast Cancer. Trends in Molecular Medicine (2017). doi:10.1016/j.molmed.2017.02.009

13. Tanaka, M. et al. Macrophage-inducible C-type lectin underlies obesity-induced adipose tissue fibrosis. Nat. Commun. (2014). doi:10.1038/ncomms5982

14. Seo, B. R. et al. Obesity-dependent changes in interstitial ECM mechanics promote breast tumorigenesis. Sci. Transl. Med. (2015). doi:10.1126/scitranslmed.3010467

15. Oudin, M. J. et al. Tumor cell-driven extracellular matrix remodeling enables haptotaxis during metastatic progression. Cancer Discov. (2016). doi:10.1158/2159-8290.CD-15-1183

16. Socovich, A. M. & Naba, A. The cancer matrisome: From comprehensive characterization to biomarker discovery. Seminars in Cell and Developmental Biology (2018). doi:10.1016/j.semcdb.2018.06.005

17. Naba, A. et al. The extracellular matrix: Tools and insights for the ‘omics’ era. Matrix Biology 49, 10–24 (2016).

18. Mayorca-Guiliani, A. E. et al. ISDoT: In situ decellularization of tissues for high-resolution imaging and proteomic analysis of native extracellular matrix. Nat. Med. (2017). doi:10.1038/nm.4352

19. Naba, A., Clauser, K. R., Lamar, J. M., Carr, S. A. & Hynes, R. O. Extracellular matrix signatures of human mammary carcinoma identify novel metastasis promoters. Elife 2014, (2014).

20. Iyengar, P. et al. Adipocyte-derived collagen VI affects early mammary tumor progression in vivo, demonstrating a critical interaction in the tumor/stroma microenvironment. J. Clin. Invest. (2005). doi:10.1172/JCI23424

21. Roussos, E. T. et al. Mena invasive (MenaINV) promotes multicellular streaming motility and transendothelial migration in a mouse model of breast cancer. J. Cell Sci. 124, 2120–2131 (2011).

22. Park, J. & Scherer, P. E. Adipocyte-derived endotrophin promotes malignant tumor progression. J. Clin. Invest. (2012). doi:10.1172/JCI63930

23. Gara, S. K. et al. Three novel collagen VI chains with high homology to the α3 chain. J. Biol. Chem. (2008). doi:10.1074/jbc.M709540200

24. Cescon, M., Gattazzo, F., Chen, P. & Bonaldo, P. Collagen VI at a glance. J. Cell Sci. (2015). doi:10.1242/jcs.169748

25. Comprehensive molecular portraits of human breast tumours. Nature 490, 61–70 (2012).

26. Price, M. A. et al. CSPG4, a potential therapeutic target, facilitates malignant progression of melanoma. Pigment Cell and Melanoma Research (2011). doi:10.1111/j.1755-148X.2011.00929.x

27. Cheng, I. H. et al. Collagen VI protects against neuronal apoptosis elicited by ultraviolet irradiation via an Akt/Phosphatidylinositol 3-kinase signaling pathway. Neuroscience (2011). doi:10.1016/j.neuroscience.2011.03.057

28. Chen, P., Cescon, M., Megighian, A. & Bonaldo, P. Collagen VI regulates peripheral nerve myelination and function. FASEB J. (2014). doi:10.1096/fj.13-239533

29. Cattaruzza, S. et al. NG2/CSPG4-collagen type VI interplays putatively involved in the microenvironmental control of tumour engraftment and local expansion. J. Mol. Cell Biol. (2013). doi:10.1093/jmcb/mjt010

30. Ojima, K., Oe, M., Nakajima, I., Muroya, S. & Nishimura, T. Dynamics of protein secretion during adipocyte differentiation. FEBS Open Bio (2016). doi:10.1002/2211-5463.12091

31. Marcelin, G., Silveira, A. L. M., Martins, L. B., Ferreira, A. V. M. & Clément, K. Deciphering the cellular interplays underlying obesityinduced adipose tissue fibrosis. Journal of Clinical Investigation (2019). doi:10.1172/JCI129192

32. Mentoor, I., Engelbrecht, A.-M., van Jaarsveld, P. J. & Nell, T. Chemoresistance: Intricate Interplay Between Breast Tumor Cells and Adipocytes in the Tumor Microenvironment. Front. Endocrinol. (Lausanne). (2018). doi:10.3389/fendo.2018.00758

33. Sherman-Baust, C. A. et al. Remodeling of the extracellular matrix through overexpression of collagen VI contributes to cisplatin resistance in ovarian cancer cells. Cancer Cell (2003). doi:10.1016/S1535-6108(03)00058-8

